# Choroid plexus tissue perfusion and secretory function in rats measured by non-invasive MRI reveal significant effects of anesthesia

**DOI:** 10.1101/2022.03.09.483707

**Authors:** Hedok Lee, Burhan Ozturk, Michael S. Stringer, Bradley J. MacIntosh, Douglas Rothman, Helene Benveniste

## Abstract

The choroid plexus (ChP) of the cerebral ventricles is a source of cerebrospinal fluid (CSF) production and also plays a key role in immune surveillance at the level of blood-to-CSF-barrier (BCSFB). In this study, we quantify ChP blood perfusion and BCSFB mediated water exchange from arterial blood into ventricular CSF using non-invasive continuous arterial spin labelling magnetic resonance imaging (CASL-MRI). Systemic administration of anti-diuretic hormone (vasopressin) was used to validate BCSFB mediated water exchange as a metric of choroidal CSF secretory function. To further investigate the coupling between ChP perfusion and CSF secretory function, we characterized the effects of two anesthetic regimens known to have large-scale differential effects on cerebral blood flow. For quantification of ChP blood perfusion a multi-compartment perfusion model was employed, and we discovered that partial volume correction improved measurement accuracy. Vasopressin significantly reduced both ChP blood perfusion and CSF secretory function. ChP perfusion was significantly higher with pure isoflurane anesthesia (2-2.5%) when compared to a balanced anesthesia with dexmedetomidine and low-dose isoflurane (1.0 %), however there was no significant difference in BCSFB water secretory function. In summary, here we introduce a non-invasive, robust, and spatially resolved in vivo imaging platform to quantify ChP perfusion as well as BCSFB water secretory function which can be applied to study coupling of these two key parameters in future clinical translational studies.

## 1. Introduction

The glymphatic system for brain waste disposal was discovered a decade ago (Benveniste et al., 2019; Iliff et al., 2012; Jessen et al., 2015; Tarasoff-Conway et al., 2015; Wardlaw et al., 2020). According to the glymphatic system model, CSF flows into peri-arterial Virchow-Robin channels and exchanges with interstitial fluid (ISF) in a manner dependent on astroglial aquaporin-4 water channels. The CSF-ISF ‘mixing’ propels waste, including amyloid beta (Aβ) and tau, towards egress routes which ultimately coalesce with authentic lymphatic vessels and the systemic circulation. Glymphatic system function and waste clearance is critically dependent on central nervous system (CNS) fluid homeostasis, including the ability to produce and reabsorb CSF at a sustainable and physiologically normal rate. It is generally assumed that 80% of CSF is continuously secreted by the choroid plexuses (ChP) (Praetorius et al., 2020) known as ‘blood to CSF barrier’ (BCSFB) function but there is a gap in knowledge of how changes in CSF production impact glymphatic waste clearance and neurodegenerative processes such as Alzheimer’s disease. This lack of information is linked to the technical challenges involved in quantifying CSF production non-invasively. For decades ChP secretory function has been quantified using the tracer dilution method which is invasive and also associated with inaccuracies (Heisey et al., 1962; Oreskovic et al., 2017). More recently an alternative invasive technique was developed which directly tracks CSF secretion from the ventricles (Karimy et al., 2015; Karimy et al., 2017). This technique recently confirmed that the CSF secretion rate in rodents depended on aging and also provided new information on the effect of various anesthetics (Liu et al., 2020). Notably, all of these methods are invasive and limits translational studies of CSF secretion in human subjects, therefore there is a great need for novel non-invasive techniques.

Recently, a non-invasive approach was introduced to quantify BCSFB mediated delivery of arterial blood water from the ChP into ventricular CSF using arterial spin labelling magnetic resonance imaging (BCSFB-ASL) (Evans et al., 2020; Perera et al., 2021). Arterial spin labelling (ASL) is a non-invasive magnetic resonance imaging technique used to quantify cerebral blood flow by using magnetically labeled inflowing arterial blood water as an endogenous tracer (Alsop et al., 2015; Grade et al., 2015; Telischak et al., 2015; Williams et al., 1992), which is increasingly used in clinical research studies of cerebral small vessel diseases, stroke, and other neurological disorders (Cada et al., 2000; De Jong et al., 1999; Ho, 2018; Iturria-Medina et al., 2016). The novel BCSFB-ASL imaging approach was previously validated by demonstrating that administration of anti-diuretic hormone vasopressin downregulated BCSFB secretory function (Evans et al., 2020). Although the new non-invasive BCSFB-ASL based approach may indeed track BCSFB secretory function, the signal to noise ratio (SNR) in ASL acquisitions is inherently poor and the BCSFB-ASL protocol requires data acquisition at long echo time (TE) that makes it even more challenging to acquire high quality and robust data. Further, quantification of ChP tissue perfusion is an important metric to quantify in parallel, to study how the coupling of ChP perfusion and secretary function changes in disease and with pharmacological manipulation. In this study we addressed these challenges by implementing a continuous ASL (CASL) sequence at 9.4 T and by applying a multi-compartment perfusion model to quantify ChP tissue perfusion as well as BCSFB secretory function.

## 2. Materials and methods

### 2.1 Animals

The animal experiments were approved by the local Institutional Animal Care and Use Committee at Yale University (New Haven, Connecticut) and conducted in accordance with the United States Public Health Service’s Policy on Humane Care. Twelve female Wistar Kyoto (WKY) rats (Charles River Laboratories, Wilmington, MA, USA) between the ages of 9 month (M) to 12M were used. All rats received standard rat chow and water ad libitum and were housed in standard conditions in a 12h light/dark cycle.

### 2.2 Anesthesia and preparation for MRI

Prior to commencing MRI acquisitions, rats underwent anesthesia induction using 3% isoflurane (ISO) delivered in 100% oxygen (O_2_). The anesthetized rats were placed in a custom 3D printed rat body holder in prone position while breathing spontaneously through a snout mask (Supplementary Material 1). For rats undergoing MRI with pure ISO (N=12), the anesthesia was maintained with 2-2.5% ISO throughout the experiments. For rats undergoing MRI under balanced anesthesia with dexmedetomidine supplemented with 1% ISO (DEXM-I) regimen (N=12), a bolus of dexmedetomidine (0.007mg/kg i.p.) was given after ISO induction and followed by continuous infusion of dexmedetomidine at a rate of 0.009 ± 0.002mg/kg/h administered via a subcutaneous catheter. In both ISO and DEXM-I experiments, ISO was delivered in a 1:1 Air:O_2_ mixture as described previously (Benveniste et al., 2017; Ozturk et al., 2021). During MRI session (protocol described below), the respiratory rate, heart rate, and body temperature were measured continuously by non-invasive MRI compatible monitors (SA Instruments, Stony Brook, NY, USA). Body temperature was maintained between 36.5–37.5°C using a heated waterbed system. Following completion of MRI scanning, the rats were recovered from anesthesia in their home cage and were observed until they regained normal behavior.

### 2.3 MRI acquisition

MRI was performed on a Bruker 9.4T/16 magnet (Bruker BioSpin, MA, USA), operating with Paravision 6.0.1 software, and interfaced with an Avance III-HD console. A custom-made transverse electromagnetic resonator style transmit-receive radiofrequency (RF) coil (Bolinger et al., 1989) with an internal diameter of 50mm with active decoupling was used for imaging (Supplementary Material 1).

#### Anatomical T2 weighted MRI

To ensure accurate and reliable positioning of the imaging slice for the CASL, two-dimensional (2D) T2-weighted (T2W) anatomical brain images were obtained in sagittal and axial orientations using a RARE sequence with the following parameters: repetition time (TR) = 2500ms, echo time (TE) = 22ms, number of acquisitions (NA) = 1, RARE factor = 4, in plane resolution = 0.23mm/voxel, slice thickness = 1.0mm.

#### Continuous arterial spin labeling

A 2D single-slice CASL sequence was performed using a single-shot spin echo EPI. Anterior commissure was identified on the anatomical T2W image and used as an anatomical reference for ensuring the inclusion of choroid plexus within the field of view; correct image plane for CASL (Supplementary Material 1). Cortical perfusion (CTX), apparent ChP perfusion, ChP tissue perfusion and BCSFB mediated water exchange were measured by incorporating a short TE followed by a long TE CASL-based acquisition. The short TE CASL sequence was performed with 9 post labelling delays (PLD = 10, 100, 200, 400, 700, 1100, 1500, 2000, 3000ms) using the following parameters: TR = 7000ms, TE = 23ms, NA = 20, acquisition matrix = 64 × 64, in plane resolution = 0.45mm/voxel, slice thickness = 1.0mm, labelling duration (LD) = 3000ms, scanning time 4min 40sec per PLD. Similarly, for the long TE CASL, the CASL sequence was performed at 9 post labelling delays (PLD = 10, 200, 400, 700, 1100, 1500, 2000, 3000, 4000ms) using following parameters: TR = 11000ms, TE=150ms, NA = 20, acquisition matrix = 64 × 64, in plane resolution = 0.45mm/voxel, slice thickness = 1.0mm, LD = 6000ms, scanning time 7min 20sec per PLD. A proton density weighted image, M_0_, was acquired by collecting control images with the following parameters: TR = 20000ms, TE = 23 and 150ms, NA = 4, scanning time 1min 20sec. A hard pulse was used for a flow-driven adiabatic inversion at 23 ± 2mm distal to the imaging plane using a separate single-turn actively decoupled arterial blood labeling coil, minimizing the magnetization transfer effect during the labeling period (Supplementary Material 1). In both the labeling and control image acquisitions, a 10mT/m labelling gradient was applied along the z direction. RF power of the labelling coil was set to 6 µT (Zhang et al., 1993). Control and labeled images were acquired in an interleaved fashion and pair-wise subtraction, *ΔM = M*_*control*_ *-M*_*labeled*_, was calculated to derive the perfusion weighted image (PWI).

#### T1 relaxometry

A 2D inversion recovery sequence was performed to calculate the tissue longitudinal relaxation time (T1) needed for perfusion measurements. A single shot and single slice echo planar imaging (EPI) imaging sequence was acquired with the same image position and spatial resolution as the CASL with the following parameters: TR = 15000ms, TE = 23ms, NA = 1, 25 inversion times ranging from 100-9000ms, acquisition matrix = 64 × 64, in plane resolution = 0.45mm/voxel, slice thickness = 1.0mm, scanning time 6min 15s.

#### T2 relaxometry

A 2D multi-spin multi-echo (MSME) sequence was performed to derive T2 distribution curves for calculating tissue and CSF partial volumes. A single slice MSME sequence was acquired with the same image position and spatial resolution as the CASL and 80 evenly spaced TEs were performed with the following parameters: TR = 6000ms, TE = 10-800ms, ΔTE = 10ms, NA = 1, acquisition matrix = 64 × 64, in plane resolution = 0.45mm/voxel, slice thickness = 1.0mm, scanning time 6min 24sec.

#### B1+ mapping

Since the accuracy of T2 multi-compartment relaxometry is sensitive to the spatially varying RF transmit (B1+) homogeneity, B1+ maps were acquired in a subset (N = 9) of the rats using a double angle method as described previously (Lee et al., 2018; Stollberger and Wach, 1996). 2D RARE sequence was performed at two flip angles in the same position and spatial resolution as the CASL using following parameters: TR = 10000ms, TE = 22ms, NA = 20, FA = 70° and 140°, RARE factor = 4, acquisition matrix = 64 × 64, in plane resolution = 0.45mm/voxel, slice thickness = 1.0mm, scanning time 11min.

### 2.4 MRI analyses

#### T1 relaxometry

Voxel-wise T1 calculations were performed by fitting image intensities of the inversion recovery sequence as a function of inversion times expressed in eq. 1 as:

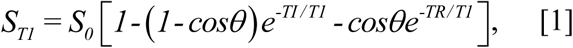

where *S*_*T1*_, *S*_*0*_, *θ, TI*, and *T1* represent image intensity, proton-density weighted signal, flip angle, inversion time, and longitudinal relaxation time, respectively. The three unknown variables (*S*_*0*_, *θ*, and *T1*) were calculated by the Nelder-Mead Simplex non-linear least square fit algorithm as described previously (Lee et al., 2018).

#### Multi compartment T2 relaxometry

Multi compartment T2 relaxometry analysis was performed to identify distinct compartments through the smooth T2 distribution curves formulated as a linear combination of exponential functions expressed in eq. 2 as

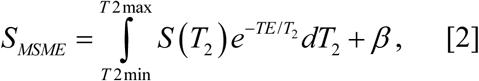

where *S(T*_*2*_) is the fractional contribution of a given T2 to the overall signal *S*_*MSME*_ and *β* is a signal offset (Does, 2018; Kroeker and Henkelman, 1986; Provencher, 1982a). *S*_*MSME*_ is defined as the mean image intensities derived from the MSME sequence within the regions of interest (CTX and ChP). The CONTIN decomposition technique (Provencher, 1982b) applies a regularized Laplace inversion and solves for an appropriate set of smooth continuous T2 distribution curve, *S(T*_*2*_), within the lower (*T2*_*min*_ *= 10ms*) and upper (*T2*_*max*_ *= 1000ms*) limits as a solution to the eq.2 (Marino, 2021; Provencher, 1982a). *dT*_*2*_ was discretized by 40 bins spaced logarithmically from *T2*_*min*_ to *T2*_*max*_. Following the calculation of a T2 distribution curve, *S(T*_*2*_) was further divided into tissue and CSF compartments whose T2 ranges were characterized by short T2 (*10ms < S(T*_*2*_*) ≤ 100ms*) in tissue and long T2 (*100ms < S(T*_*2*_*) ≤ 1000ms*) in CSF and the areas under the distribution curves within the limits defined the fractional contribution of the tissue (*P*_*tissue*_) and CSF (*P*_*CSF*_) compartments. An expectation value, *E[T*_*2*_], whose probability density function was defined by the T2 distribution curve, was calculated to identify the peak of T2 within the range of each compartment.

#### Evaluation of CONTIN algorithm using computer simulations

The accuracy of the CONTIN decomposition technique for calculating the T2 distribution curve was evaluated using computer simulations incorporating a broad range of measurement conditions. MSME image intensity, *S*_*simulated*_, was simulated by a bi-exponential function expressed in eq. 3 as

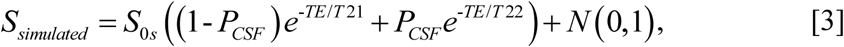

where *S*_*0s*_, *T21, T22, and N(0,1)* represent the simulated water proton signal, T2 of tissue at 40ms, T2 of CSF at 300ms, and a noise term defined by a normal distribution with a fixed mean (0) and standard deviation (1), respectively. *S*_*simulated*_ was calculated at 80 evenly spaced TEs from 10ms to 800ms and *S*_*0s*_ was set to 60, 100, and 200 in simulating various SNR conditions. A total of 5000 trials was performed for a given set of variables (*S*_*0s*_, *0.1 ≤ P*_*CSF*_ *≤ 0.9*) and *E[T*_*21*_*], E[T*_*22*_], and *P*_*CSF*_ were calculated at each trial. Discrepancy between the calculated and the given set of values (*T*_*21*_ = 40ms *T*_*22*_ = 300ms and *P*_*CSF*_ = 0.1 - 0.9) were quantified and expressed in eq. 4 as

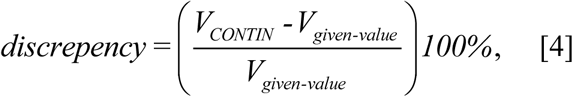

where *V*_*CONTIN*_ and *V*_*given-value*,_ represent the calculated and corresponding given values in *E[T*_*21*_*], E[T*_*22*_], and *P*_*CSF*_, respectably.

#### Manual delineations of regions of interest

ChP and somatosensory CTX were manually delineated as shown in Fig.1. Perfusion weighted images taken at TE = 23ms, *ΔM*_*TE=23ms*_, were summed over all PLDs and used as a template for identifying the CTX. Similarly, at the level of the ChP, the BCSFB weighted images acquired at TE=150ms, *ΔM*_*TE=150ms*_, were summed over all PLDs and the conspicuous regions within the lateral ventricles were delineated. A publicly available free software MRICron (https://www.nitrc.org/projects/mricron) was utilized for the manual delineations.

**Figure. 1.**
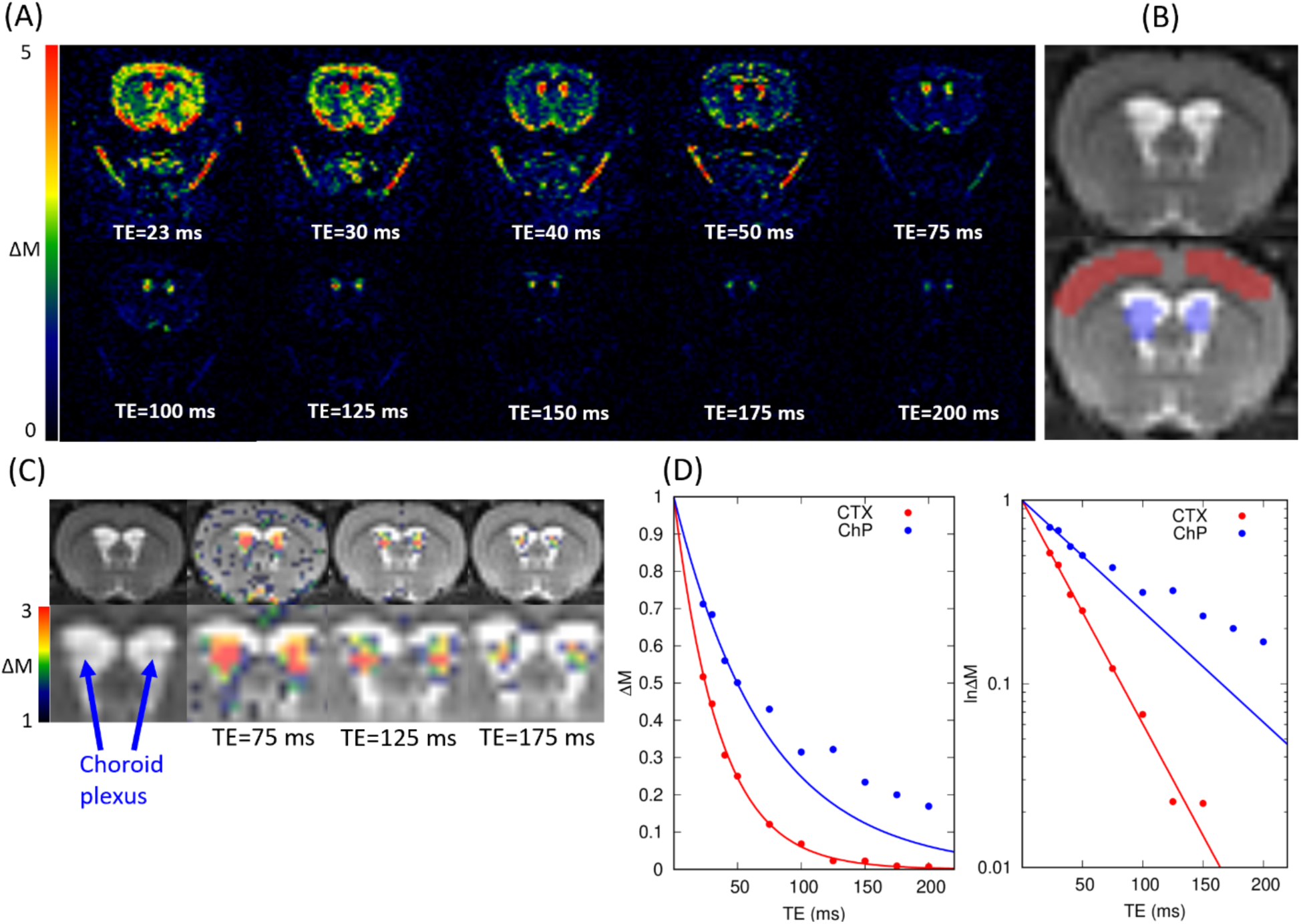
(A) Representative perfusion weighted images, *ΔM*, in colormaps are shown as a function of echo times. Images were taken using following CASL parameters: TR = 11000ms, LD = 6000ms, and PLD = 500ms (B) T2W anatomical scan and corresponding regions of interest in cortex (red) and choroid plexus (blue). (C) Perfusion weighted images taken at TE= 75, 125 and 175 ms (in color) are overlaid onto the corresponding T2W anatomical scan (in grey scale). (D) Perfusion weighted signals (N = 2) within the regions of interest are plotted as a function of echo times in cortex (red) and choroid plexus (blue). The solid lines represent the mono-exponential curve fittings using the TE = 23, 30, 40, 50ms. Both linear (left) and semi-logarithmic (right) plots are shown.

#### Cortical perfusion, apparent ChP perfusion, and BCSFB measured by CASL

In the short TE CASL measurement, the coarse spatial resolution of the CASL sequence relative to the size of the ChP manifests as signal mixing between the ChP tissue perfusion signal (*f*_*ChP*_) and the BCSFB water exchange signal (*f*_*BCSFB*_), analogous to the partial volume effect between different tissue compartments (Asllani et al., 2008; Chappell et al., 2018; Chappell et al., 2021). The signal mixing resulting in apparent ChP perfusion, *f*_*ChP-BCSFB*_, was modeled as the sum of the partial contributions of *f*_*ChP*_ and *f*_*BCSFB*_ expressed in eq. 5 as

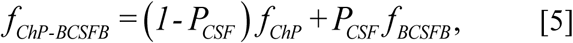

where *f*_*ChP-BCSFB*_ and *f*_*BCSFB*_ were calculated directly from the CASL measurements taken at short TE and long TE CASL, respectively, as expressed in eq. 6 and *P*_*CSF*_ was calculated from the T2 relaxometry.

The calculations of CTX perfusion (*f*_*CTX*_), apparent ChP perfusion (*f*_*ChP-BCSFB*_), and BCSFB (*f*_*BCSFB*_) were performed by implementing a standard single compartment model (Alsop et al., 2015; Buxton et al., 1998), where *f*_*CTX*_ and *f*_*BCSFB*_ represent the rate of labelled arterial blood water exchanging with tissue and CSF, respectively. In the single compartment model, the relationship between the pair-wise subtraction, *ΔM=M*_*control*_ *-M*_*labeled*_, and *f*_*CTX*_ and *f*_*BCSFB*_ was expressed by

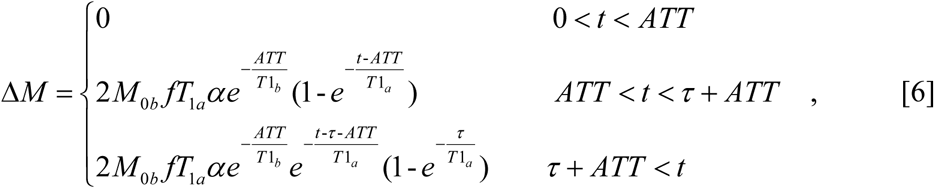

where, *f, T*_*1b*_, *α, ATT*, and *τ* represent the tissue perfusion (or BCSFB) given in the units of ml/min/100g, T1 of blood assumed to be 2400ms, the arterial blood inversion efficiency assumed to be 0.8, the arterial transit time, and the arterial blood labeling duration, respectively (Dobre et al., 2007; Hirschler et al., 2020). Note that *t* is the time measured from the start of the arterial blood water labeling to the time of the image acquisition (*LD+PLD*). *f* and *ATT*, in eq. 6, were solved by fitting *ΔM/M*_*0b*_ as a function of PLDs using the Levenberg–Marquardt non-linear least square algorithm written in-house MATLAB code (MathWorks, Natick, MA, USA). Initial conditions of *f* and *ATT* were set to 200 and 500ms, respectively, and the parameter search range was constrained within following limits: 10ml/100g/min ≤ *f* ≤ 500ml/100g/min and 10ms ≤ *ATT* ≤ 3000ms (Larkin et al., 2019; Thomas et al., 2006). *f*_*CTX*_ and *f*_*ChP-BCSFB*_ were calculated by using the CASL sequence acquired at TE = 23ms (*ΔM*_*TE=23 ms*_) and *f*_*BCSFB*_ was calculated by using the CASL sequence acquired at TE = 150ms (*ΔM*_*TE=150 ms*_). *M*_*0b*_ in both *f*_*CTX*_ and *f*_*ChP-BCSFB*_ represents the equilibrium magnetization of arterial blood which was approximated by the equilibrium magnetization of tissue, *M*_*0-TE=23ms*_, derived from the image acquired at TR = 20000ms and TE = 23 ms which was then divided by the water tissue to blood partition coefficient, assumed to be 0.98 in *f*_*CTX*_ and 1.05 in *f*_*ChP-BCSFB*_ (Chappell et al., 2018; Herscovitch and Raichle, 1985). *T*_*1a*_ in *f*_*CTX*_ and *f*_*ChP-BCSFB*_ was approximated by an apparent T1 of CTX and ChP measured by the inversion recovery T1 mapping technique taken at TE = 23ms. In *f*_*BCSFB*,_ *M*_*0b*_ was approximated by the equilibrium magnetization of CSF, *M*_*0-CSF*_, however *M*_*0-CSF*_ cannot be directly measured since the ROI of ChP comprises partial volume effect of ChP tissue and CSF, therefore *M*_*0-CSF*_ was estimated by using the linear regression algorithm which extrapolated the image intensity of ChP taken at TR = 20000ms and TE = 150ms, *M*_*0-TE=150ms*_, at the limit of pure CSF (Supplementary Material 2) (Asllani et al., 2008; Parkes et al., 2004).

#### Effects of vasopressin on ChP perfusion and BCSFB secretory function

To characterize the effects of the anti-diuretic hormone vasopressin, a subset of female WKY rats (N=8) were used. Following anesthesia with 2-3% ISO delivered in 100% oxygen and with the rats breathing spontaneously, a PE-50 femoral infusion line was placed for intravenous administration of vasopressin (Vasostrict Par Pharmaceutical Companies, USA). The rats were subsequently transported to the MRI scanner with the intravenous femoral infusion line long enough to continuously infuse vasopressin from the outside of the scanner using a micro-infusion pump (Legato 130 syringe pump, KD Scientific, USA). During imaging the rats were anesthetized with 2-3% ISO delivered with a 1:1 Air:O_2_ mixture and breathing spontaneously. The pharmacological MRI (ph-MRI) protocol comprised short (23ms) and long (150ms) TE CASL at a fixed PLD acquired in an interleaved manner (Perera et al., 2021) using the following parameters TR = 11000ms, TE = 23/150ms, number of acquisitions (NA) = 10/20, acquisition matrix = 64 × 64, in plane resolution = 0.45mm/voxel, slice thickness = 1.0mm, LD = 6000ms, PLD = 500/2000ms. Following baseline scans, comprising three blocks of short and long TE scans, a continuous infusion of vasopressin was commenced at a rate of 2mU/Kg/min (Faraci et al., 1990). Post-administration scans included a total of six blocks of short and long TE CASL scans lasting for 60minutes. Semi-quantitative image analysis was performed by calculating the % signal change between the average of the three baseline scans and the post administration scans.

#### Effects of anesthesia on ChP perfusion and BCSFB secretory function

The modulatory effect of the two anesthetics regimens (DEXM-I and ISO) on *f*_*CTX*_, *f*_*ChP*_, *f*_*ChP-BCSFB*_, and *f*_*BCSFB*_, were carried out using a crossover repeated measures study design as described previously (Ozturk et al., 2021). All rats (N = 12) underwent two MRI scanning sessions in which the rats were randomly selected to undergo either DEXM-I or ISO in the first scanning session. Before the second scanning session, at least 7 days of rest was allowed to ensure wash-out effects of each anesthetic. In the second session, all rats underwent either DEXM-I or ISO so that all rats received both anesthetics over the two scanning sessions totaling 24 imaging sessions. The data are presented as mean estimates for the CASL outcomes with standard deviation (sd). For experiments with repeated measurements on each animal, pairwise comparisons were investigated using Wilcoxon signed-rank 2-sided test. Analyses were performed using the SPSS (New York, USA) statistical software package and a p-value less than 0.05 was considered significant. Linear regression analyses were performed between the apparent ChP perfusion and ChP tissue perfusion with respect to BCSFB secretory function using XLSTAT statistical data analysis software (New York, USA)

## 3. Results

### 3.1 TE-dependent perfusion weighted signal decay

Representative perfusion weighted images, *ΔM*, are shown in Fig.1A as a function of TEs. As seen from the figure, the signal from the brain tissue reached background noise levels around TE = 150ms, however, the signals within the lateral ventricles (LV), remained conspicuous even at TE = 200ms, implying an inherently longer T2 relaxation time compared to the brain tissue. Notably, the spatial distribution of *ΔM* image intensity within the LV was heterogenous as can be observed in Fig.1C. Specifically, while *ΔM* ‘hot spots’ were observed in the center of the LV, higher signals were associated with the ChP itself. Therefore, the source of hot spot regions in the center of the LV taken at TE =150ms or longer is BCSFB mediated water exchange in the CSF compartment of ChP.

Perfusion weighted signals in the CTX and ChP were calculated and plotted as a function of TEs (Fig.1D). Consistent with the observations in Fig.1A, CTX signals decayed considerably faster than ChP signals and in the CTX the mono-exponential characterization of *ΔM* fitted well even at the long TEs (red solid line, Fig.1D). In contrast, the mono-exponential curve fitted the ChP poorly in the end portion of the curve (TE = 100ms or greater), implying the existence of multiple compartments in the region of interest.

### 3.2 Evaluation of the CONTIN algorithm by computer simulations

The accuracy of the CONTIN algorithm for calculating T2 distribution curves in the simulated bi-exponential model, as expressed in eq.3, was evaluated by computer simulation modeling. At each trial, simulated bi-exponential data was analyzed by the CONTIN algorithm and a calculated T2 distribution curve was analyzed to derive, *E[T*_*21*_*], E[T*_*22*_], and *P*_*CSF*_ and compared with the given values (*T*_*21*_ = 40ms *T*_*22*_ = 300ms and *P*_*CSF*_ = 0.1 - 0.9). In Fig.2A averaged T2 distribution curves of all 5000 trials with *P*_*CSF*_ = 0.6 are shown at three different SNRs: 60, 100, 200. As expected, bimodal T2 distribution curves with two peaks at 40ms and 300ms were discernable. Line broadening around the peaks is caused by the noise in the simulations and becomes more pronounced with decreasing SNR but the broadening did not cause the peaks to overlap, and two peaks remained easily distinguishable. Discrepancies between the calculated and the given values *E[T*_*21*_*], E[T*_*22*_] and *P*_*CSF*_ are shown in the Fig.2B-D. Notably, with increasing SNR the accuracy improved in all of the three variables as expected and the accuracy of the calculated variables are dependent on the given fractional contribution, *P*_*CSF*_. With increasing *P*_*CSF*_, the peak at *T*_*22*_ = 300ms appeared more distinct, improving the detectability and reducing the discrepancies in calculating *T*_*22*_ and *P*_*CSF*_. Conversely with decreasing *P*_*CSF*_, the discrepancies in calculating the variables amplified.

**Figure. 2.**
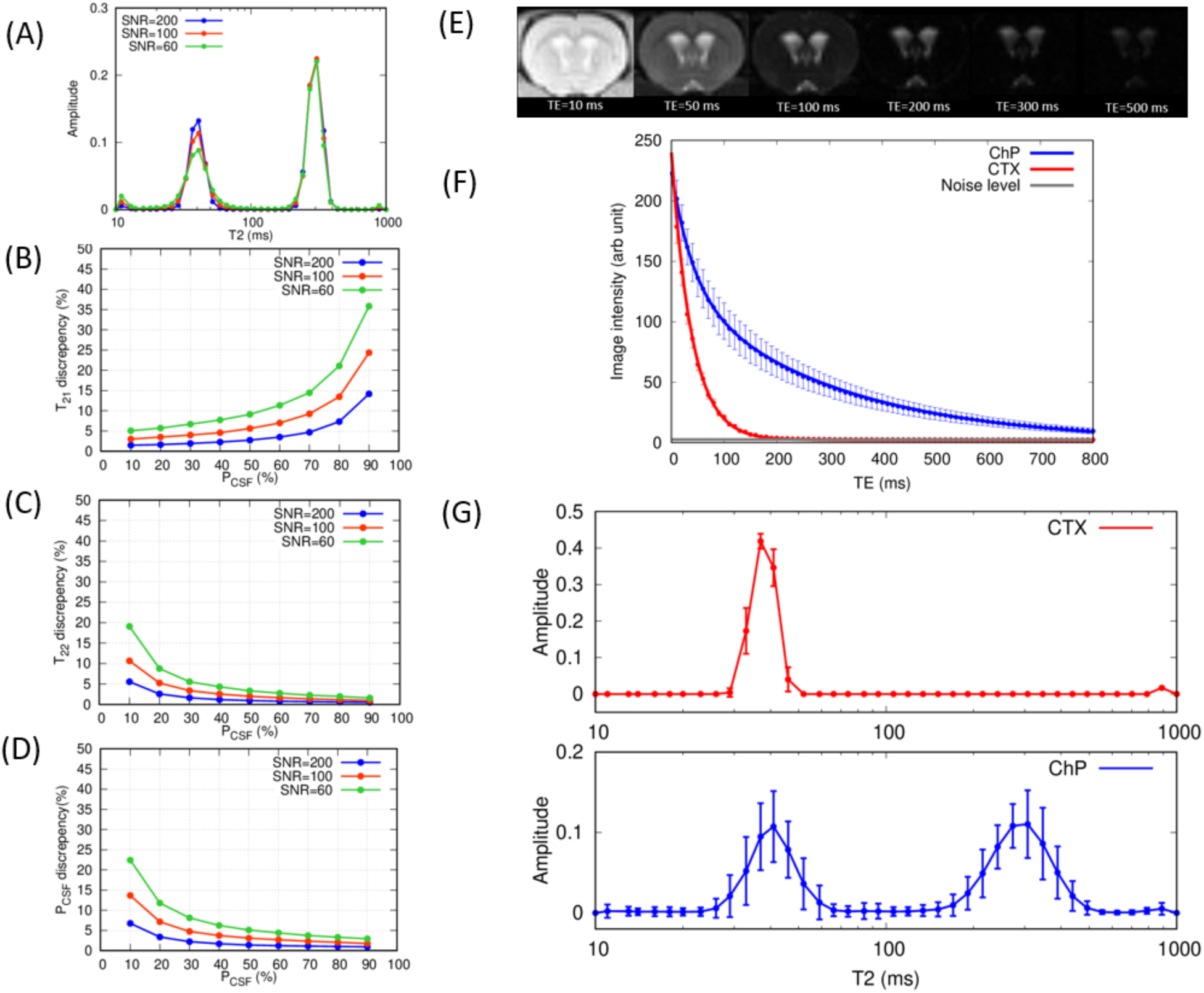
(A) Averaged T2 relaxation distribution curves calculated by the CONTIN algorithm in the computer simulations when *P*_*CSF*_ = 0.6 at SNR = 60, 100, 200. (B)(C)(D) Discrepancies between the calculated and given values are plotted as a function of *P*_*CSF*_ for (B) *T*_*21*_ (C) *T*_*22*_ and (D) *P*_*CSF*_. (E) Representative images acquired in MSME sequence at TE = 10, 50, 100, 200, 300, 500ms. (F) Population averaged (mean ± sd N=24) MSME signals are plotted as a function of echo times in cortex (red) and choroid plexus (blue). Solid lines represent the multi-compartment T2 relaxometry fits. (G) Population averaged (mean ± sd N=24) T2 relaxation distributions curves in cortex (red) and choroid plexus (blue) are shown. DEXM-I (N=12) and ISO (N=12) data were combined in the results in (F) and (G).

### 3.3 T2 relaxometry in choroid plexus and cortex in rats

For each animal, the mean image intensities within the ROIs were calculated over all TEs and the population averaged mean intensities are plotted in Fig.2F along with the corresponding representataive images (Fig.2E). Similar to the trend observed in the perfusion weighted images (Fig.1), the spin-echo image intensities also decayed more rapidly in CTX than ChP and reached background noise levels around TE = 150ms while the intensities associated with the ChP decayed more gradually and retained signal above the noise level even at TE = 800ms. At TE = 10ms, the SNR of the ChP was 103 ± 7 and the SNR of the CTX was 91 ± 6, derived by using a canonical form for calculating SNR (Firbank et al., 1999). Population averaged T2 distribution curves are also shown in Fig.2G. The T2 distribution curve in CTX manifested a unimodal distribution curve peaking at *E[T*_*21*_] = 38 ± 1ms implying a single compartment. In contrast, a bimodal distribution curve was observed within the ROI of the ChP with short and long T2 peaks at *E[T*_*21*_] = 41 ± 4ms and *E[T*_*22*_] = 301 ± 16ms, respectively. Coarse MR image resolution relative to the size of ChP, as shown in Fig.1C, inevitably results in partial volumes across ChP tissue and the surrounding CSF and the fractional contribution of CSF compartment, *P*_*CSF*_, was 0.57 ± 0.8. Finally, the RF transmit inhomogeneity (B1+) was also analyzed within the ROI of ChP as it can significantly result in stimulated echoes corrupting the measured MSME signals (Prasloski et al., 2012). Nine rats underwent B1+ mapping using the double angle method (Stollberger and Wach, 1996) and yielded 0.953 ± 0.012 indicative of a highly homogeneous B1+ profile owing to the use of the volume transmit coil along with the ChP being positioned near the center of the coil.

### 3.4 Vasopressin MRI

Pharmacodynamics of the anti-diuretic hormone vasopressin was characterized through the interleaved CASL sequence acquired at short (23ms) and long (150ms) TEs with fixed PLDs. Representative time averaged (3 frames) perfusion and BCSFB weighted images are shown in Fig.3A. Notably, the perfusion weighted image intensities appear to increase progressively in the CTX after administration of vasopressin. In contrast, both the apparent ChP perfusion and the BCSFB weighted image intensities decreased progressively over time with vasopressin administration. Semi-quantitative characterization of the effect of vasopressin was assessed by calculating % signal change between the pre-(baseline) and post-vasopressin data for all animals and the results are plotted in Fig.3B. As seen from the representative images, the perfusion in CTX gradually increased during vasopressin administration and stabilized ∼25% above the baseline after 60min, while both the apparent perfusion and BCSFB weighted signals of the ChP gradually decreased over time, reaching ∼25% below the baseline at 60min. In spite of the progressive decrease of the BCSFB weighted signals over time, implying downregulation of BCSFB function, the volume of the lateral ventricles neither shrunk nor expanded as shown in Fig.3A.

**Figure. 3.**
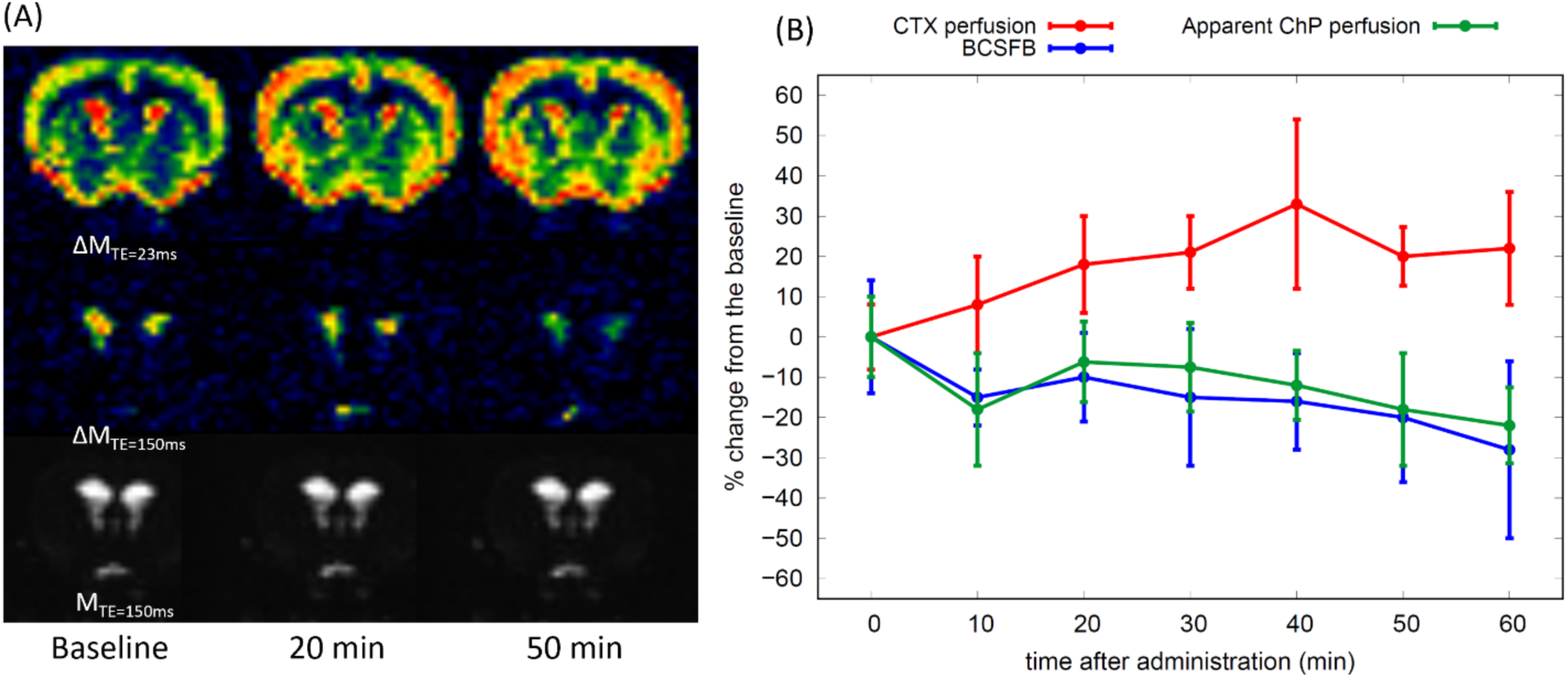
(A) Representative perfusion weighted (*ΔM*_*TE=23 ms*_), BCSFB weighted (*ΔM*_*TE=150 ms*_), and control (*M*_*TE=150ms*_) images are shown over the three time points. Images were taken at TR = 11000ms, TE = 23ms, 150ms, LD = 6000ms, PLD = 500ms. (B) % signal change (mean ± sd) between the pre (baseline) and post administration of vasopressin (N=8) are shown as a function of time.

### 3.5 Effects of anesthesia on choroid plexus perfusion and BCSFB

Normalized ChP perfusion and BCSFB weighted signals, (*ΔM*/*M*_*0b*_), were fitted using the single compartment model expressed in eq.6 for deriving CTX perfusion, the apparent ChP perfusion, and BCSFB and the results are shown in Fig.4A-C and summarized in Table.1. *ΔM*/*M*_*0b*_ in all the animals (N = 12) are plotted as dots along with the group mean ± sd and the solid lines represent the corresponding model fitting using the population averaged *f* and *ATT* plotted separately for the two anesthetic regimens of DEXM-I and ISO.

**Table 1.**
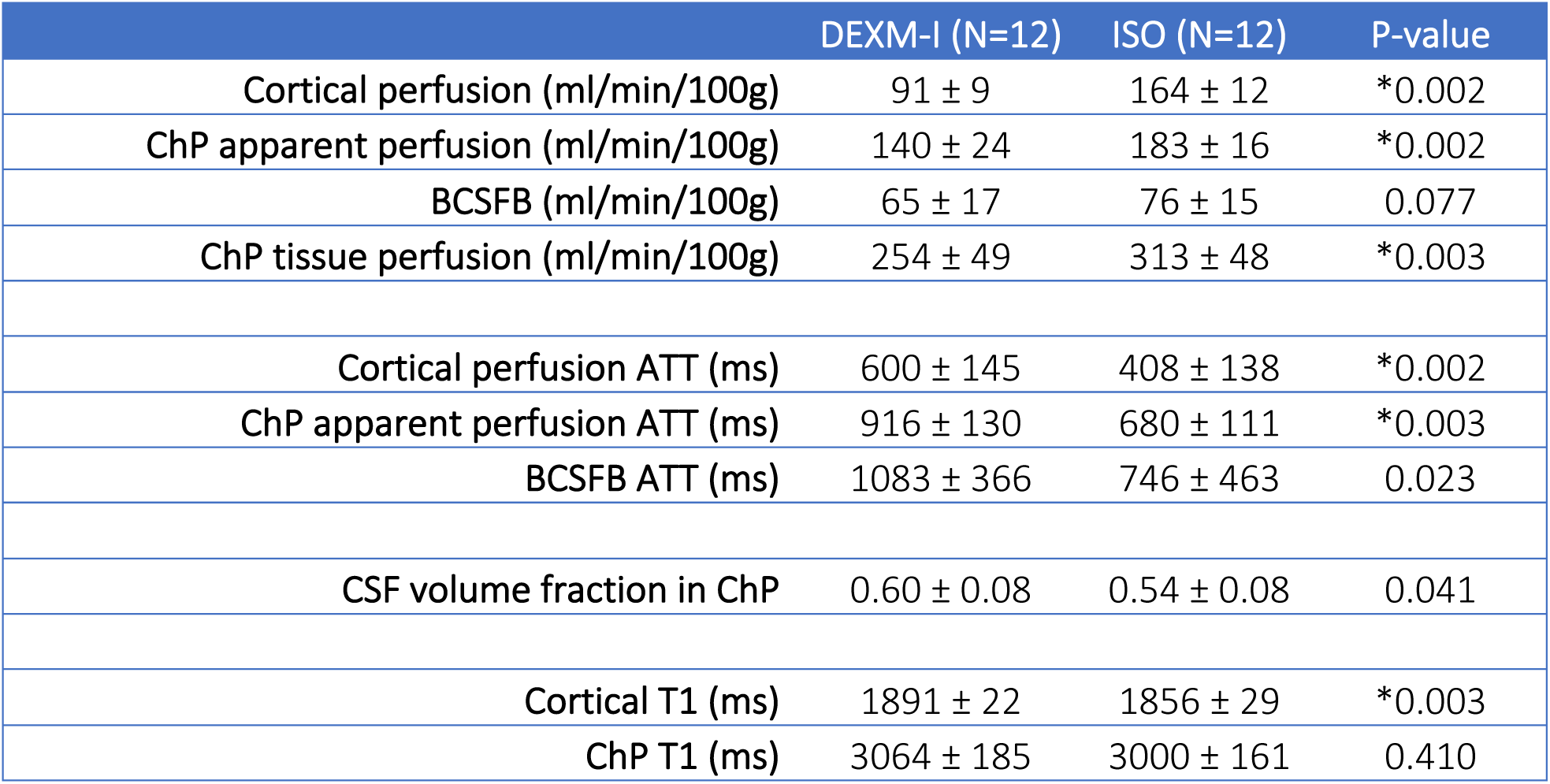
Results of perfusion and BCSFB measurements between DEXM-I versus ISO. A summary of variables calculated in the single and multi-compartment perfusion models, multi-compartment T2 relaxometry and T1 relaxometry are given as mean ± sd. p-value is calculated by Wilcoxon signed-rank 2-sided test. *P<0.05 is considered significant.

**Figure. 4.**
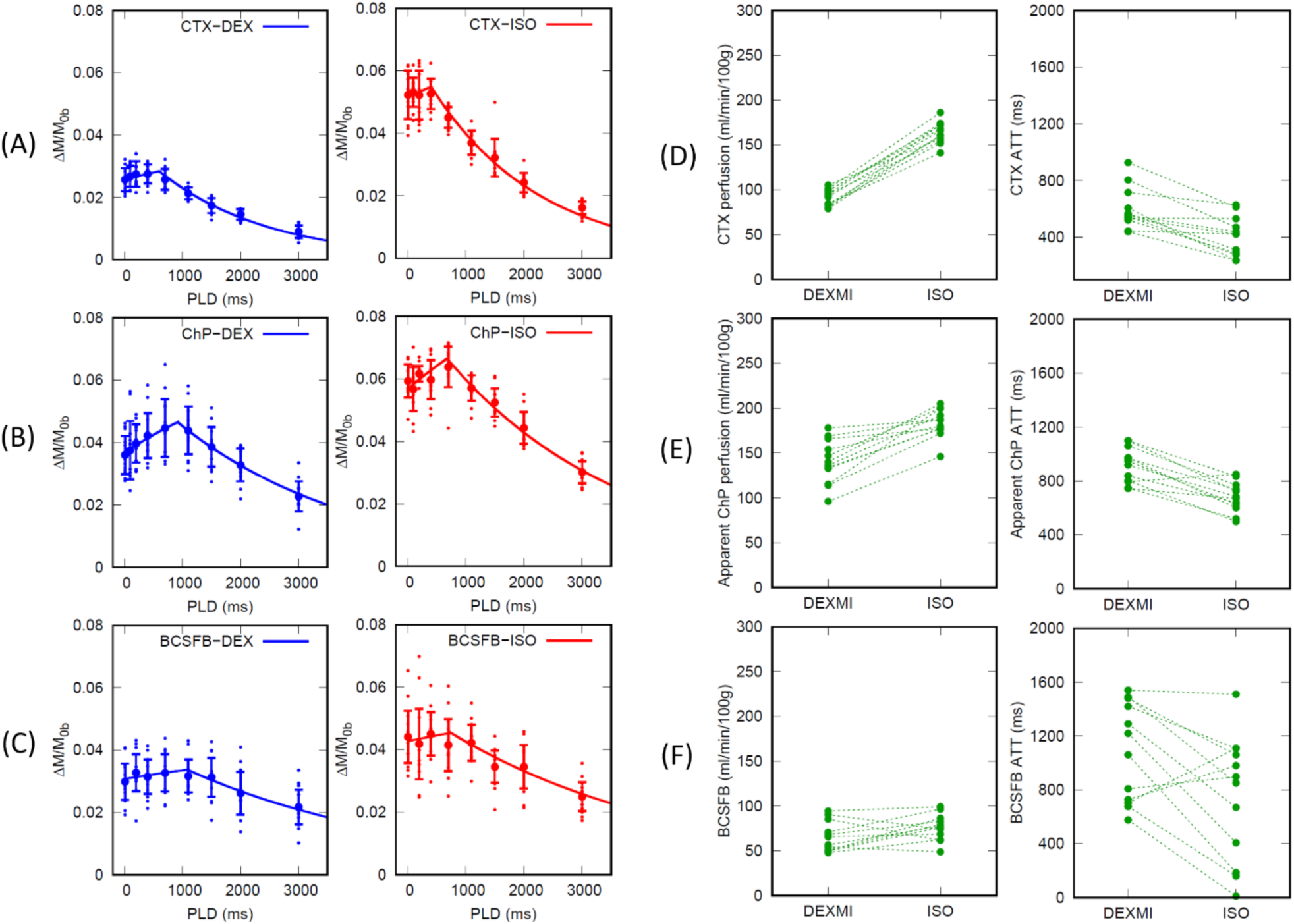
(A) (B) Normalized perfusion weighted signals (mean ± sd) in cortex (CTX) and choroid plexus (ChP) are plotted as a function of PLDs in all animals (N = 12). Solid lines represent the single compartment model fits using the population averaged perfusion and ATT. DEXM-I (blue) and ISO (red) results are plotted. (C) Normalized BCSFB weighted signals (mean ± sd) within choroid plexus are plotted as a function of PLDs in DEXM-I (blue) and ISO (red). Solid lines represent the single compartment model fits using the population averaged BCSFB and ATT. Ladder plots represent (D) CTX perfusion, (E) apparent ChP perfusion, and (F) BCSFB in each animal.

As summarized in Table.1 and the ladder plots in Fig.4D-F ChP tissue perfusion was two to three folds higher than the CTX perfusion and the BCSFB was only 25% of the ChP tissue perfusion values. The duration for the arterial blood to flow from the labeling region to the tissue, ATT, was also significantly longer in the apparent ChP perfusion compared to CTX irrespective of the anesthetic regimen. In addition, the duration for the arterial blood to arrive at the CSF through the BCSFB mediated water exchange was nearly doubled compared to that of the CTX perfusion. A Wilcoxon signed-rank 2-sided test was performed to compare the effect of the two anesthetics and yielded significant differences in the apparent ChP perfusion, ChP tissue perfusion, and CTX perfusion. ATTs were also significantly lower with ISO compared to DEXM-I in the apparent ChP perfusion and CTX perfusion likely due to the vasodilatory effect of ISO, elevating blood flow, thereby shortening the duration for the arterial blood to flow from the labeling region to reach the tissue (Wegener et al., 2007). Spatially resolved post labelling delay (PLD) summed perfusion and BCSFB weighted images are shown in Fig.5A and 5B. Reduced brain tissue perfusion with DEXM-I anesthesia compared to ISO is strikingly noticeable. As for the BCSFB weighted images, prominent signals of labelled blood water exchanging with CSF in the LV are conspicuous but differences between the two distinct anesthetic regimens are not discernable and statistical analyses of BCSFB values between the two anesthetics yielded a 13% reduction with DEXM-I compared to ISO but was not statistically significant (p-value = 0.077). Linear regression analysis revealed significant correlations between the apparent ChP perfusion and BCSFB secretion (p-value = 0.012) and between ChP tissue perfusion and BCSFB (p-value = 0.020), as shown in the Fig.5C and D, suggesting that blood to CSF barrier mediated delivery of arterial blood water from the ChP into ventricular CSF is directly proportional to the ChP tissue perfusion, however, the rate of exchange is not one to one. Note also that low intensity signals in the BCSFB weighted images, as shown in the Fig 5B, were also observed in various areas of the CSF compartment including the LVs, cortical surface, and peri-vascular spaces of the basal arteries in agreement with data reported in recent study in humans, and potentially indicative of labelled blood water exchanging with CSF (Petitclerc et al., 2021).

**Figure. 5.**
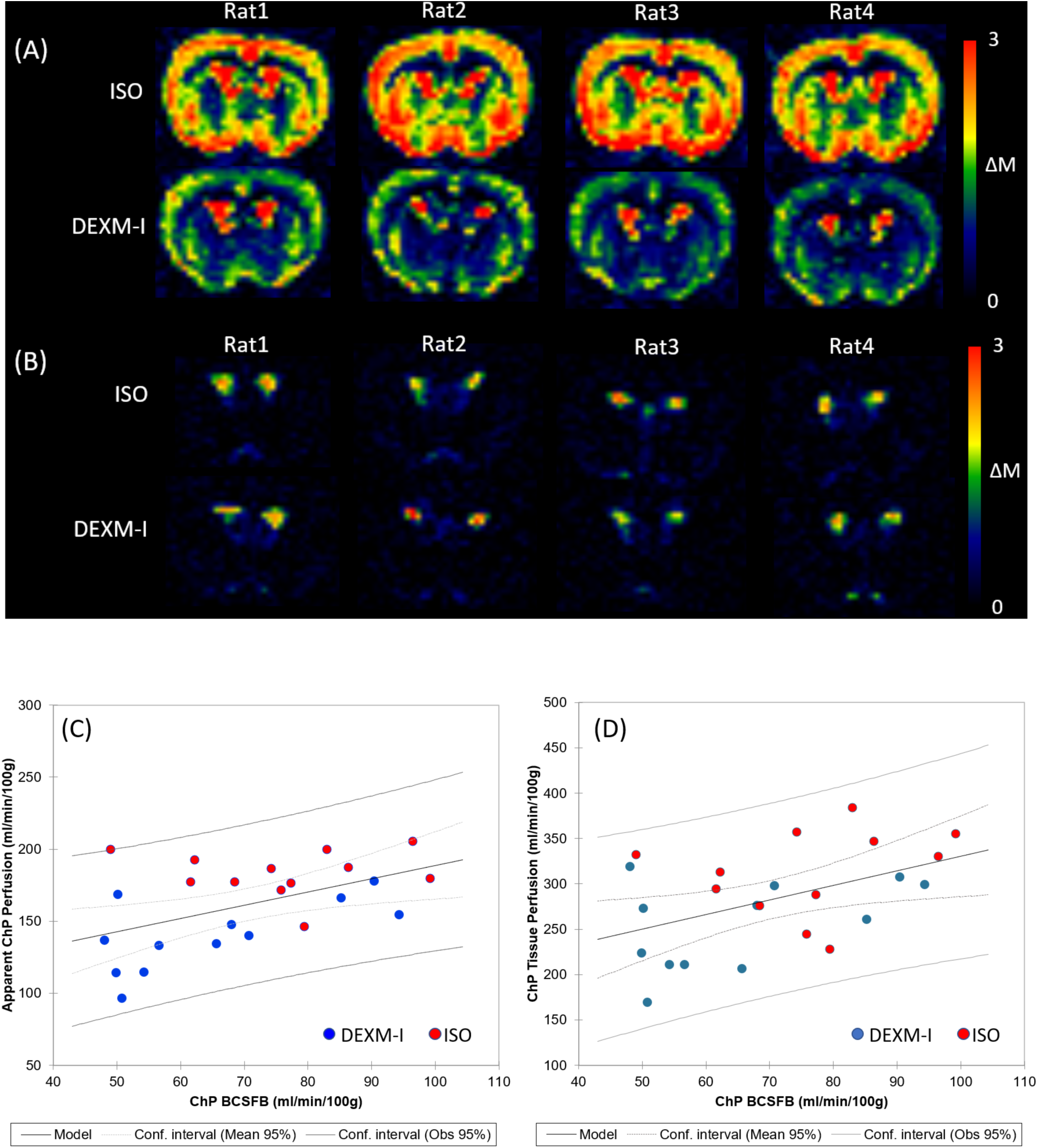
(A) Perfusion weighted images (*ΔM*_*TE=23ms*_) summed over all PLDs are shown in 4 rats undergoing ISO or DEXM-I anesthesia. (B) BCSFB weighted images (*ΔM*_*TE=150ms*_) summed over all PLDs are shown in 4 rats undergoing ISO or DEXM-I anesthesia. (C) Apparent choroid plexus perfusion and (D) choroid plexus tissue perfusion are plotted as a function of BCSFB. Red and blue dots represent 12 rats undergoing ISO and DEXM-I anesthesia, respectively. Solid and dotted lines represent the linear regression lines and confidence intervals, respectively.

## 4. Discussion

The objective of this study was to quantify ChP tissue perfusion and ‘arterial blood water exchange with CSF’, also known as ‘BCSFB’ using the non-invasive CASL MRI imaging technique. BCSFB-ASL is an attractive non-invasive method for studying BCSFB function as it measures the unidirectional water movement from the ChP to the CSF compartment. In our study we implemented CASL which use a long pulse labelling duration to enhance SNR permitting acquisition of data at higher spatial resolution than previous studies in mice which used a pulsed ASL sequence (Evans et al., 2020; Perera et al., 2021). From our CASL experiments we draw the following main conclusions: (1) the ChP perfusion-weighted signals acquired at TE = 23ms volume average across ChP tissue perfusion as well as the BCSFB signals; (2) when the CASL protocol was applied to suppress the tissue perfusion signals to near background noise (acquisitions at TE = 150ms), the BCSFB-ASL signal within the LV was spatially heterogeneous and the BCSFB-ASL ‘hot spot’ signals coincided with the exact locations of ChP tissue; (3) we validated the BCSFB-ASL sequence in the rat brain by confirming downregulation of ChP perfusion as well as BCSFB secretory function with systemic administration of vasopressin (Evans et al., 2020; Perera et al., 2021), and (4) we observed differential effects of two anesthetic regimens on the coupling of ChP perfusion and BCSFB function.

We developed a novel processing approach and modelled the ChP perfusion-weighted signal taken at TE = 23ms as the sum of the partial signal contributions from ChP tissue perfusion and BCSFB secretion. While the CTX perfusion-weighted signals plotted as a function of TE adhered to a mono-exponential signal decay, the perfusion-weighted signal decay in the ChP deviated from the mono-exponential decay at later TEs, implying signal mixing between multiple compartments. Interestingly, other MRI modalities such as diffusion and T2 relaxometry studies share similar attributes that involve complex interactions between water and the macro and micro-environment of the cellular and interstitial matrix, which result in multi-exponential signal decays with respect to parameters like TEs and b-factors (Alexander et al., 2001; Does, 2018; Graham et al., 1996; Niendorf et al., 1996; Riemer et al., 2018). In this study using CASL we uncovered that the ChP perfusion-weighted signal did not follow a mono-exponential decay due to partial volume effects of ChP tissue perfusion and BCSFB function. Further, without correcting for the partial volume effects the ChP tissue perfusion was significantly underestimated (nearly 2-to 3-fold) underscoring the importance of considering partial volume effects when studying ChP tissue perfusion, at least in a small animal species such as rodents. However, it remains an open question whether similar partial volume effects will also impact ChP tissue perfusion measurement in humans and more studies are needed to clarify this important issue (Alisch et al., 2021; Johnson et al., 2021; Petitclerc et al., 2021; Zhao et al., 2020).

Partial volume effects in ASL have been studied extensively in human cerebral perfusion experiments and often employ computerized methods to segment each voxel into the three tissue compartments: grey matter (GM), white matter (WM) and CSF (Chappell et al., 2021; Zhao et al., 2020). Due to the lack of publicly available software and algorithms to reliably segment the ChP into distinct compartments in rodents, the present study employed T2 relaxometry which allowed us to probe distinct compartments without a priori assumptions about the number of compartments within the ChP (Does, 2018; Graham et al., 1996; Serradas Duarte and Shemesh, 2018). Here we used the smoothed T2 distribution curves to identify the different compartments. Specifically, one compartment was observed in the CTX (unimodal peak) and two compartments in the ChP (bimodal peaks).

A single peak observed in CTX at T2 ∼40ms was consistent with a previous study and overlapped with a peak of a shorter T2 compartment within the ChP (de Graaf et al., 2006). We therefore concluded that the ChP tissue and CTX share identical T2s and that the apparent T2 in the ChP is significantly overestimated without the abovementioned partial volume effect correction (Alisch et al., 2021). The other peak noticed in the ‘ChP’ was characterized by a long T2 ∼300ms which, we identified as CSF since no other tissue or blood compartment retains such long T2 (Daoust et al., 2017; de Graaf et al., 2006). Note that numerical Laplace transformation is known to be sensitive to noise and imposes relatively stringent requirements on experimental conditions in order to attain accurate and robust results (Does, 2018; Graham et al., 1996). Therefore, we evaluated the CONTIN algorithm using simulated bi-exponential signals with noise levels identical to our experiment. The results indicated that the derived parameters were within a 5% margin of error suggesting that our experimental conditions are sufficient to accurately derive T2 distribution curves and fractional volume calculations.

Whether or not the rate of water exchange from blood into CSF, termed BCSFB, is directly related with CSF secretory function in the ChP still needs further investigation. To further validate this claim, we performed simultaneous recordings of CTX perfusion, the apparent ChP perfusion and BCSFB using an interleaved CASL acquisition with short- and long TE CASL, and also evaluated the pharmacological effect of vasopressin, a drug known to reduce ChP perfusion and consequently secretory function. Vasopressin – also called anti-diuretic hormone - interacts with at least 2 types of receptors, V1 and V2, and is known to regulate brain water homeostasis to mediate its physiological effects through V1 receptors situated on the ChP epithelial cells.

Significant reduction in ChP perfusion by vasopressin in a dose dependent manner has previously been reported and attributed to stimulation of V1 receptors in the smooth muscle cells of choroidal blood vessels causing vasoconstriction and increased vascular resistance (Brinton et al., 1984; Faraci et al., 1988; Fernandez et al., 2001; Phillips et al., 1988; Vakili et al., 2005). Another study reported that if vasopressin is administered as a continuous infusion the ChP perfusion as well as the CSF secretion is significantly reduced (Faraci et al., 1994). The study by Faraci et al., strongly implied that ChP perfusion and CSF secretory function were tightly coupled and downregulated with systemic administration of vasopressin. Our vasopressin results are in agreement with previous studies and support the BCSFB metric as a surrogate marker for CSF secretory function (Evans et al., 2020; Perera et al., 2021). In addition to the downregulation of BCSFB by vasopressin, we also noted that the CTX perfusion progressively increased consistent with previous reports (Chung et al., 2003; Kozniewska and Szczepanska-Sadowska, 1990; Perera et al., 2021; Suzuki et al., 1993). It has been suggested that the mechanism underlying the vasopressin-induced increase in cerebral perfusion is mediated by V2 receptors or the release of nitric oxide from the endothelium (Chung et al., 2003; Suzuki et al., 1993). However, here we offer an alternative explanation. Inhalational anesthetics, like isoflurane used in relatively high concentrations (∼2-2.5%), is known to partially obliterate cerebral autoregulation which under normal conditions keeps the cerebral perfusion constant and within a normal physiological range of mean arterial pressure (Gropper, 2019). We used isoflurane in the vasopressin experiments and we contribute the observed effect of vasopressin – a vasoconstrictor elevating the blood pressure – as a consequence of ISO-induced attenuation of autoregulation and the cerebral perfusion consequently becoming dependent on arterial pressure (Chung et al., 2003; Kozniewska and Szczepanska-Sadowska, 1990). As always, care should be taken in interpreting the results of cerebral hemodynamics and BCSFB function in anesthetized animals (Perera et al., 2021).

We also tested the effect of two different anesthetics and observed that the cerebral perfusion was nearly 50% lower in the DEXM-I group compared to the ISO group and these differences appeared to be global. Dexmedetomidine is a potent and selective agonist of the alpha-2 adrenergic receptor known to induce cerebral vasoconstriction and lower CBF in mice and humans and our data are consistent with these data (Petrinovic et al., 2016; Prielipp et al., 2002; Zornow et al., 1990; Zornow et al., 1993). The effects of anesthetics on BCSFB function is of particular interest for studies of glymphatic transport because the choice of anesthesia and brain arousal state can profoundly affect solute transport but scarce data exists on the coupling between brain waste clearance and CSF secretory function (Benveniste et al., 2017; Hablitz et al., 2019; Xue et al., 2020). The choroid plexus is densely innervated by sympathetic nerve fibers and its secretory and hemodynamic functions are sensitive to sympathetic nerve stimulation or denervation (Johanson et al., 2008; Lindvall et al., 1978; Lindvall and Owman, 1981; Liu et al., 2020; Oreskovic et al., 2017). At the molecular level, known drivers for CSF secretory function in the ChP has been attributed to osmotic gradients between CSF and blood serum and AQP1 water channels (Praetorius et al., 2020). A recent study in mice using the direct CSF production measurement technique reported 33% higher CSF formation with isoflurane compared to a balanced anesthesia with an α2-adrenergic agonists such as with Ketamine-Xylazine (Liu et al., 2020). Based on these data we expected DEXM-I to reduce BCSFB function compared to ISO but our data did not corroborate this hypothesis. In a previous study, ISO anesthesia was reported to sustain CSF production but affecting its reabsorption (Artru, 1984a, b). In our study we documented a statistically significant coupling of ChP perfusion and secretory function but reports also vary in this regard. For example, no clear correlation between ChP perfusion and rate of CSF secretory function was reported during hypercapnia (Oppelt et al., 1963), or with wide variations of PaCO_2_ and hypotension (Carey and Vela, 1974; Martins et al., 1976; Weiss and Wertman, 1978). These discrepancies are likely due to the commonly used tracer dilution method for measurement of CSF secretion, which is known to be sensitive to the experimental conditions (Ivan et al., 1971). More studies using BCSFB-ASL are needed to evaluate the sensitivity and reliability of this technique compared to existing invasive methods and possibly establish standardized protocols similar to cerebral perfusion ASL measurements (Alsop et al., 2015).

A number of limitations related to the imaging and analysis methods used in our study need to be considered. First, the manual delineation of the ChP inevitably might have introduced CSF contamination into the perfusion weighted data, and to circumvent such potential errors a voxel-wise perfusion calculation can instead be performed similar to a recent human study (Petitclerc et al., 2021). However, increased spatial resolution would introduce more noise into the BCSFB signals and other means to improve SNR in the data acquisitions should be considered (e.g., cryogenically cooled RF receiver coils (Li et al., 2021; Stanton et al., 2021). Second, the calculation of the T2 distribution curves by the CONTIN algorithm and the associated computational processing time and linewidths of the T2 distribution curves can be further improved using other fitting methods like non-negative least square algorithm (Graham et al., 1996). Third, the spatial variations of B1+ over the imaging volume in the MSME sequence resulted in imperfect refocusing with stimulated echoes thereby potentially corrupting the data (Weigel, 2015). Thus the accuracy of T2 distribution curves can be further improved by considering the B1+ inhomogeneities in a curve fitting algorithm (Serradas Duarte and Shemesh, 2018). Fourth, our model excluded the CASL signal contribution of the intravascular compartment of ChP. Although bipolar gradients were employed it may have been insufficient to completely null the vascular contribution, thereby potentially overestimating the apparent ChP perfusion. Therefore, a more complex three compartment model to explicitly model intravascular contribution with intercompartmental exchange may improve the accuracy of ChP tissue perfusion measurements (Gregori et al., 2013; Petitclerc et al., 2021; Wells et al., 2013). Lastly, while BCSFB secretory function tended to be higher with ISO compared to DEXM-I the effect did not reach conventional statistical significance, potentially due to the relatively small sample size (N=12) which may have lacked sufficient power to capture subtle effects as revealed in the linear relationship between ChP tissue perfusion and BCSFB.

## Credit authorship contribution statement

Hedok Lee: Conceptualization, Methodology, Software, Investigation, Writing. Burhan Ozturk : Methodology, Investigation. Michael Stringer: Conceptualization, Writing. Bradley MacIntosh: Conceptualization, Writing. Douglas Rothman: Methodology, Writing. Helene Benveniste: Conceptualization, Methodology, Supervision, Funding acquisition, Writing.

## Acknowledgments

The author(s) disclosed receipt of the following financial support for the research, authorship, and/or publication of this article: The present work was supported by National Institutes of Health R01AT011419 (H.B.), Foundation Leducq Transatlantic Network of Excellence (16/CVD/05) (H.B.) and PureTech Health (H.B.).

The authors thank Peter Brown and Terence Nixon of MRRC (Magnetic Resonance Research Center) at Yale University for coil development and support.

## Data and declaration of conflicting interests

The author(s) declared no potential conflicts of interest with respect to the research, authorship, and/or publication of this article.

## Supplementary materials

Supplemental materials for this article are available online.

## Data availability statement

Imaging data and analysis software for the present study are available through reasonable request to the corresponding author.

## Notes

### Competing Interest Statement

The authors have declared no competing interest.

