## Supplementary for "Choroid plexus tissue perfusion and secretory function in rats measured by non-invasive MRI reveal significant effects of anesthesia"

### Supplementary material 1

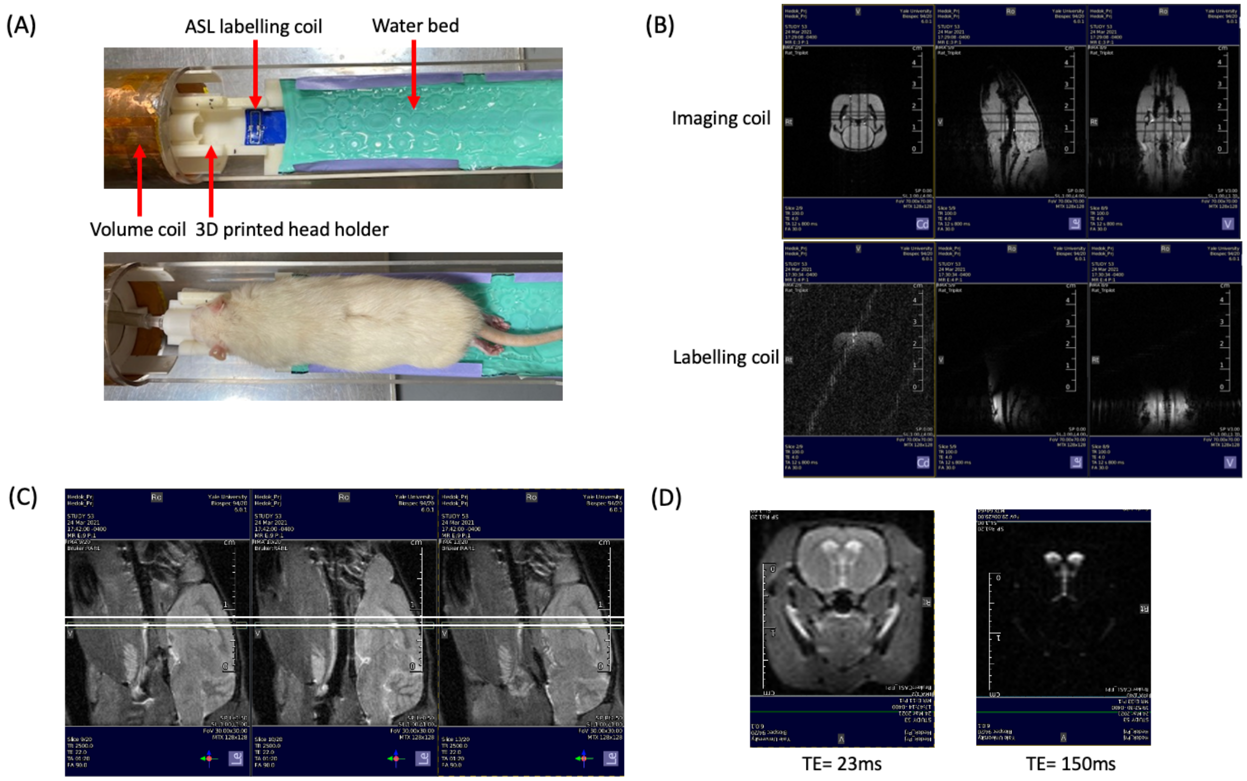


Supplementary Figure.1

(A) A 3D printed animal holder to place rat in a prone position, volume transmit and receive coil (ID = 50mm), and actively decoupled ASL labelling coil are shown. (B) Anatomical localizer scans captured by the imaging and labelling coils along three orthogonal planes. (C) T2W anatomical scan captured along sagittal plane. Solid lines represent an ASL image plane. (D) Representative untagged CASL single shot EPI image taken at TE = 23ms and 150ms.

**Supplementary material 2**

$M_{0b}$ in BCSFB was calculated by dividing the $M_{0\text{-}CSF}$ by the CSF–blood partition coefficient 1.15 (Chappell et al., 2018). A kernel based regression algorithm was implemented to estimate $M_{0\text{-}CSF}$ using $M_{0\text{-}TE=150ms}$ (Asllani et al., 2008; Chappell et al., 2021). The lateral ventricle, which was manually delineated as shown in the Supplementary Figure.2A, defined the perimeter of kernel comprising the broad range of partial volume between choroid plexus tissue and CSF within the region of interest. The partial volume was modeled as a linear function expressed as,

$$M_{0\text{-}TE=150ms} = \alpha \, T1_a + \beta \qquad [s1]$$

$$M_{0\text{-}CSF} = M_{0\text{-}TE=150ms} \quad \text{when } T1_a = 4000ms \qquad [s2]$$

where $T1_a$ represents the longitudinal relaxation time measured at TE = 23ms. By sampling $M_{0\text{-}TE=150ms}$ and $T1_a$ in each voxel within the perimeter and $\alpha$ and $\beta$ were calculated using a linear least square algorithm. $M_{0\text{-}CSF}$ was then estimated at the limit of pure CSF when $T1_a$ =4000ms as shown in the Supplementary Figure.2B.

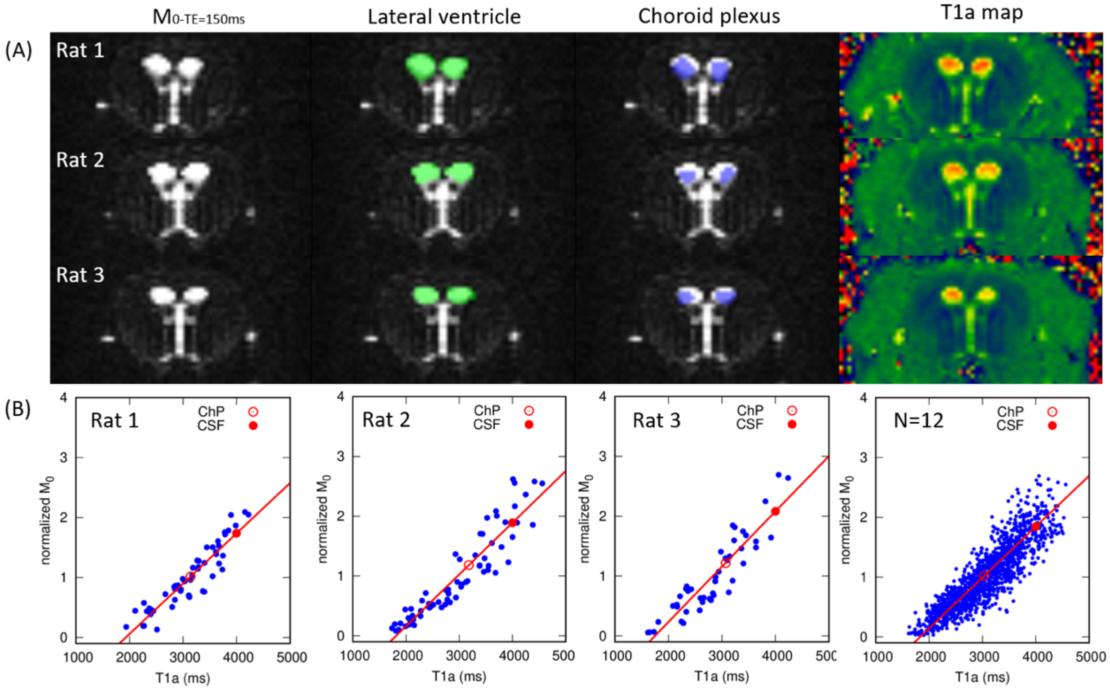


Supplementary Figure.2

(A) Manually delineated regions of interest (ROIs) are overlaid onto the corresponding $M_{0\text{-}TE=150ms}$ image in three animals: lateral ventricle (green) choroid plexus (blue). Corresponding T1 maps taken at TE = 23ms (T1a) are also shown. (B) Normalized $M_0$ ($M_{0\text{-}TE=150ms}$ /mean intensity within LV) image intensities are plotted as a function of T1a within the lateral ventricle. Normalized $M_0$ at ChP (T1a ~ 3000ms) and CSF (T1a = 4000ms) are shown in open and filled circles, respectively. Solid lines represent the linear regression lines as expressed in eq. s1.
